# Dynamic label-free analysis of SARS-CoV-2 infection reveals virus-induced subcellular remodeling

**DOI:** 10.1101/2023.11.16.567378

**Authors:** Nell Saunders, Blandine Monel, Nadège Cayet, Lorenzo Archetti, Hugo Moreno, Alexandre Jeanne, Agathe Marguier, Timothy Wai, Olivier Schwartz, Mathieu Fréchin

**Author notes:** Contributed equally (Lorenzo Archetti, Hugo Moreno, Alexandre Jeanne).

## Abstract

Assessing the impact of SARS-CoV-2 on organelle dynamics allows a better understanding of the mechanisms of viral replication. We combine label-free holo-tomographic microscopy (HTM) with Artificial Intelligence (AI) to visualize and quantify the subcellular changes triggered by SARS-CoV-2 infection. We study the dynamics of shape, position and dry mass of nucleoli, nuclei, lipid droplets (LD) and mitochondria within hundreds of single cells from early infection to syncytia formation and death. SARS-CoV-2 infection enlarges nucleoli, perturbs LD, changes mitochondrial shape and dry mass, and separates LD from mitochondria. We then used Bayesian statistics on organelle dry mass states to define organelle cross-regulation (OCR) networks and report modifications of OCR that are triggered by infection and syncytia formation. Our work highlights the subcellular remodeling induced by SARS-CoV-2 infection and provides a new AI-enhanced, label-free methodology to study in real-time the dynamics of cell populations and their content.

## Introduction

The COVID-19 pandemic is caused by the severe acute respiratory syndrome-coronavirus-2 (SARS-CoV-2)^1^, inducing a broad spectrum of syndromes from a light cold to life-threatening pneumonia^2^. The search for SARS-CoV-2 treatments is continuing^3^ and reductionist approaches vastly dominate experimental efforts. A stop-motion view of the SARS-CoV-2 infection cycle has emerged^4^ where its impact on the host cell is understood through key host/virus molecular entanglements^5^. Previous studies tackled the impact of the virus on a global cellular scale, employing fluorescence and electron microscopy^6,7^ and as such were lacking the dimension of time. Filming the impact of SARS-CoV-2 on an entire cellular system from early infection to death would greatly improve our understanding of infection sequences and dynamics, yet the efforts to obtain such knowledge are precluded by the limitations of live microscopy. The various types of fluorescence microscopy induce non-neglectable phototoxicity and molecular perturbations due to the use of chemical or genetic labeling^8–13^. This limits the capacity to observe multiple targets over hours-long periods, which is the time scale necessary to capture the cellular changes induced by SARS-CoV-2. Classical label-free imaging techniques such as phase contrast or differential interference contrast (DIC), while less invasive, provide images plagued by optical aberrations, poor contrast, and limited spatial resolution. A new generation of AI-augmented label-free microscopic methods has emerged, emulating fluorescence staining for key cellular structures in the absence of fluorescence^14,15^, or bolstering the usage of lower-content, label-free images to detect specific cellular states^16–18^. Holo-tomographic microscopy (HTM) provides high-content refractive index (RI) images able to capture complex biological processes and multiple cellular structures at unprecedented spatial resolution and ultralow-power illumination^19^. When combined with computer vision, HTM can support image-based quantitative investigations of cell dynamics over hours at relevant temporal resolutions^19^.

In this study, we developed a high-content imaging pipeline combining live HTM, machine learning and Bayesian statistics to provide a quantitative and dynamic vision of the impact of SARS-CoV-2 on the organelle system of hundreds of infected cells in culture.

## Results

### Label-free microscopy shows virus-induced cellular alterations

Through key host-viral protein interactions, SARS-CoV-2 reshapes the subcellular organization and the organelles of its target cells^6,7^. SARS-CoV-2 reroutes lipid metabolism^20–22^, fragments the Golgi apparatus^7^, promotes the formation of double-membrane vesicles^7,23,24^ and alters mitochondrial function^25^, with the goal of boosting virus production while delaying antiviral responses^26,27^. Our aim was to capture the kinetics and extent of such alterations in living cells by recording and quantifying cellular and organellar dynamics in real time using HTM^19^. We selected U2OS-ACE2 cells as targets because of their high sensitivity to SARS-CoV-2 and their flat shape that facilitates imaging^28^. Cells were first infected with the Wuhan strain and imaged with HTM (Figure 1A and seen <u>here</u>). Time lapse experiments were carried for up to two days or until the death of infected cells (Figure 1B). Non-infected cells were recorded as a control. The most obvious event visible as soon as 10 hours post infection (pi) was the formation of syncytia, a known phenomenon where infected cells expressing the viral spike (S) protein at their surface fuse with neighboring cells^28,29^.

**Figure 1.**
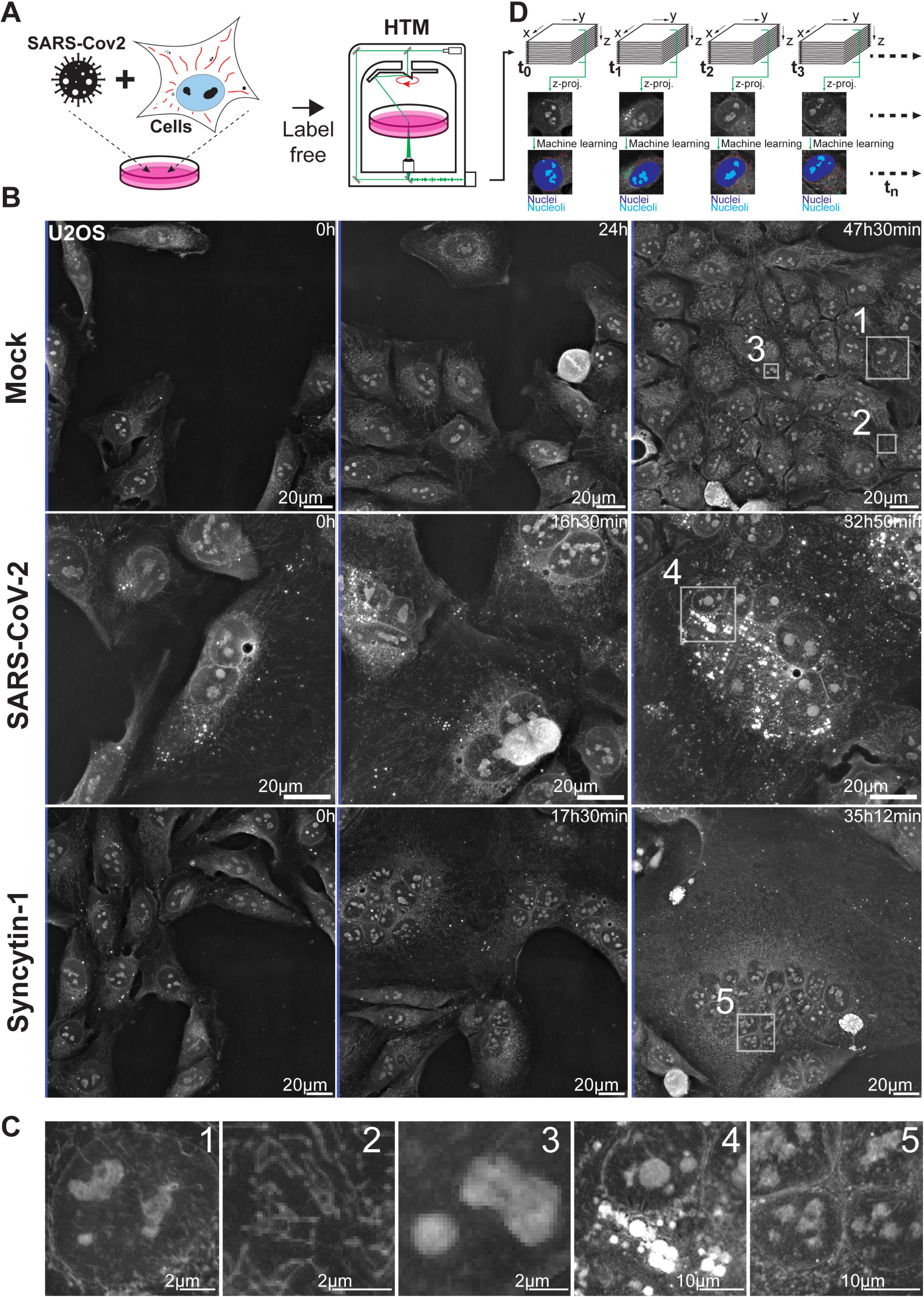
SARS-Cov-2 infected cells show subcellular dynamic changes. (**A**) Refractive index (RI) map of U2OS cells infected with SARS-Cov-2, acquired using holotomographic microscopy (HTM). (**B**) Representative images of non-infected cells (Mock), SARS-CoV-2 infected cells or cells forming syncytia upon expression of the fusogenic protein Syncytin-1 (**C**) Magnifications of cellular details available for further ML-aided image analysis, such as nuclei (1), mitochondria (2), nucleoli (3), lipid droplets (4) or syncytia nuclei cluster (5). (**D**) Time lapse imaging data projection for 2D computer-vision analysis and qualitative assessment of cells.

Formation of syncytia was used as a marker of productively infected cells. In such cells, we noticed a quick clustering of nuclei, visible as soon as two or more cells started to fuse. The zone of nuclei clustering apparently hosted groups of growing lipid droplets (LD), accumulating over time, while mitochondria were moving away from this region and redistributed across the cytoplasm. Within the nuclei, nucleoli appeared denser and rounder upon infection (Figure 1C). We next determined which of these cellular events were due to the infection itself or the result of syncytia formation. To differentiate between these possibilities, we recorded cells that fused together in the absence of SARS-CoV-2, after transient expression of Syncytin-1, a fusogenic protein involved in the formation of placental syncytiotrophoblasts^30^. Infection-independent syncytia did not show the same features. LD remained small and rare, and nucleoli were not altered (Figure 1B and 1C). However, we detected similar mitochondrial movements in both SARS-CoV-2- and Syncytin-1-induced syncytia. As demonstrated before^19^, we did not detect the Golgi network, the cytoskeleton, nor DMVs with HTM since these structures show little RI contrast with their surroundings.

To go beyond the qualitative nature of these observations and to quantify the cellular alterations triggered by SARS-CoV-2, we designed a HTM image quantification pipeline where cells and organelles were detected in time-lapse recordings by tailored machine learning (ML) approaches (Figure 1D).

### Machine learning detects cellular organelles in high-resolution label-free images

We adapted our ML strategy to the different characteristics of the biological objects of interest. Mitochondria that are small, pixel-scale objects, were detected using a two-class pixel categorization^31^ where a trained extra-tree classifier attributes a class to each pixel, based on its position in a derived feature space. Such an approach allowed precise and accurate detection of these sparse objects within the highly textured HTM images (Figure 2A). Its large hyperparameter space was not explored through a *human in the loop* process but using the optuna optimization framework^32^. Larger objects such as nuclei, nucleoli and whole cells were optimally segmented with an adapted U-NET^33^ fully convolutional network (Figure 2B). For whole cell segmentation, a sharpening of the outlines was performed by object propagation within the RI signal^34^. LD were segmented using a Nanolive assay which automatically detects LD based on their high refractive index, unique signal distribution and roundness.

**Figure 2.**
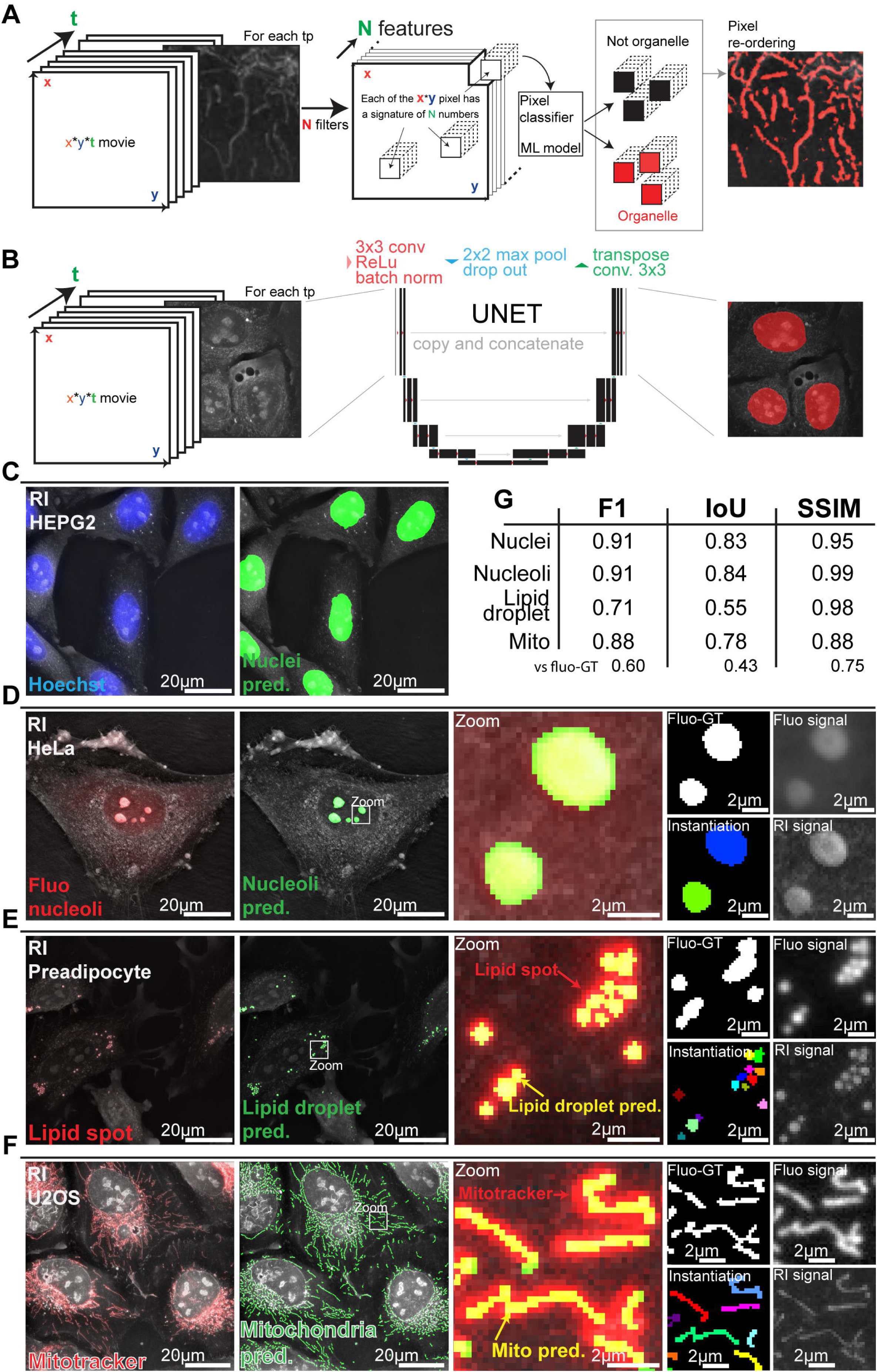
Machine learning detects key organelles within HTM images. **(A)**Mitochondria and lipid droplets detection using pixel classification: a feature space of size x*y*N is calculated by applying N convolution filters on each image time point (tp) of size x*y. An extra tree classifier decides for each pixel if it belongs or not to an organelle signal based on its position in the feature space. **(B)** Nuclei, nucleoli, and cells detection using the convolutional network UNET. **(C-F)** Comparison of **(C)** nuclei, **(D)** nucleoli, **(E)** lipid droplet and **(F)** mitochondria detection within refractive index images with their respective fluorescent label signal. Structures are thicker when visualized with epifluorescence microscopy compared to holotomography. **(G)** F1 score, intersection over union (IoU) score and structural similarity index measure (SSIM) for each organelle. Nuclei, nucleoli and lipid droplet prediction scores are calculated against fluo-derived ground truths. Mitochondria are evaluated against fluo-derived ground truth and expert hand labeling of mitochondria within the refractive index image.

We then validated the automatic segmentations of organelles within RI images by labelling the cells with organelle-specific fluorescent markers. Nuclei, nucleoli, LD, and mitochondria were stained respectively with Hoechst, Green Nucleolar staining (ab139475), lipid spot, and Mitotracker DeepRed (Figure 2C-2G). We used the standard F1 and intersection over union (IoU) scores for the strict binary evaluation of masks versus references, and the structural similarity index measure (SSIM)^35^ for a quantification of similarity perception. For nuclei and nucleoli that are large and simple objects, matches between masks and fluorescent signals were high (Figure 2C and 2G). The LD masks were not perfectly matching with the lipid spot fluorescent signal. This was expected, since HTM resolving power is better than epifluorescence^19^ (Figure 2E). This illustrates the challenge of objectively quantifying the quality of few-pixels object masks, especially in a live context where biological structures move through the succession of acquisition regimes. For these reasons, the scores of LD predictions were very good yet slightly lower than those of nuclei and nucleoli (Figure 2G).

Similarly, our RI-based mitochondrial predictions were sharper and better resolved than the fluorescent signal generated by Mitotracker DeepRed (Figure 2F). The scores obtained from comparing our RI-based ML predictions against fluorescence-derived references were good (Figure 2G) yet lower than those of the other organelles because of unavoidable mismatches between predictions and ground truth. In addition to motion, the typical crowding of mitochondria in the perinuclear region generates unresolved^36^ Mitotracker epifluorescence signal. Such signal is not optimal for comparison purposes. We thus used an expert-generated segmentation of mitochondria within a RI image to assess further the quality of our ML-generated mitochondrial mask (Figure 2G).

To validate the biological relevance of our mitochondrial detection workflow, we silenced *OPA1*, a dynamin-like GTPase protein required for mitochondrial fusion^37^ whose ablation causes mitochondrial fragmentation and inherited optic neuropathy^38^. Inspection of the label-free HTM images revealed an obvious fragmentation of the mitochondrial network and our ML-based mitochondrial detection system reported a reduction of mitochondria size distribution (Figure S1 and S2). This confirmed our capacity to automatically detect and quantify mitochondrial morphology under basal and pathological conditions in a label-free manner.

### SARS-CoV-2 infected cells display specific organelle dynamics

We then examined in real-time the subcellular changes induced by infection with two different SARS-CoV-2 strains. We recorded movies of U2OS cells infected with either the SARS-CoV-2 Wuhan ancestral virus or the Omicron BA.1 variant. As controls, we used uninfected cells and cells that underwent intercellular fusion upon Syncytin-1 expression. We used our algorithms to follow individual or fused cells and to segment nuclei, nucleoli, LD, mitochondria and the cytosol, the latter being defined by the subtraction of all other compartments from the cell. This analysis was performed every 12 minutes, from 2 to 48 h pi. Our object segmentation was robust, allowing analysis over time within each movie, and across movies (Figure 3 and Video S1-19). Of note, the formation of syncytia did not alter the quality of our predictions. Nuclei and nucleoli were properly detected despite the appearance of compact clusters of nuclei in syncytia (Figure 3A and 3B). LD remained well-defined even when their size increased or when they moved from the cytosol to the perinuclear region (Figure 3C and 3D). Mitochondria were accurately segmented, despite a large variety in length and distribution in infected cells (Figure 3C and 3D).

**Figure 3.**
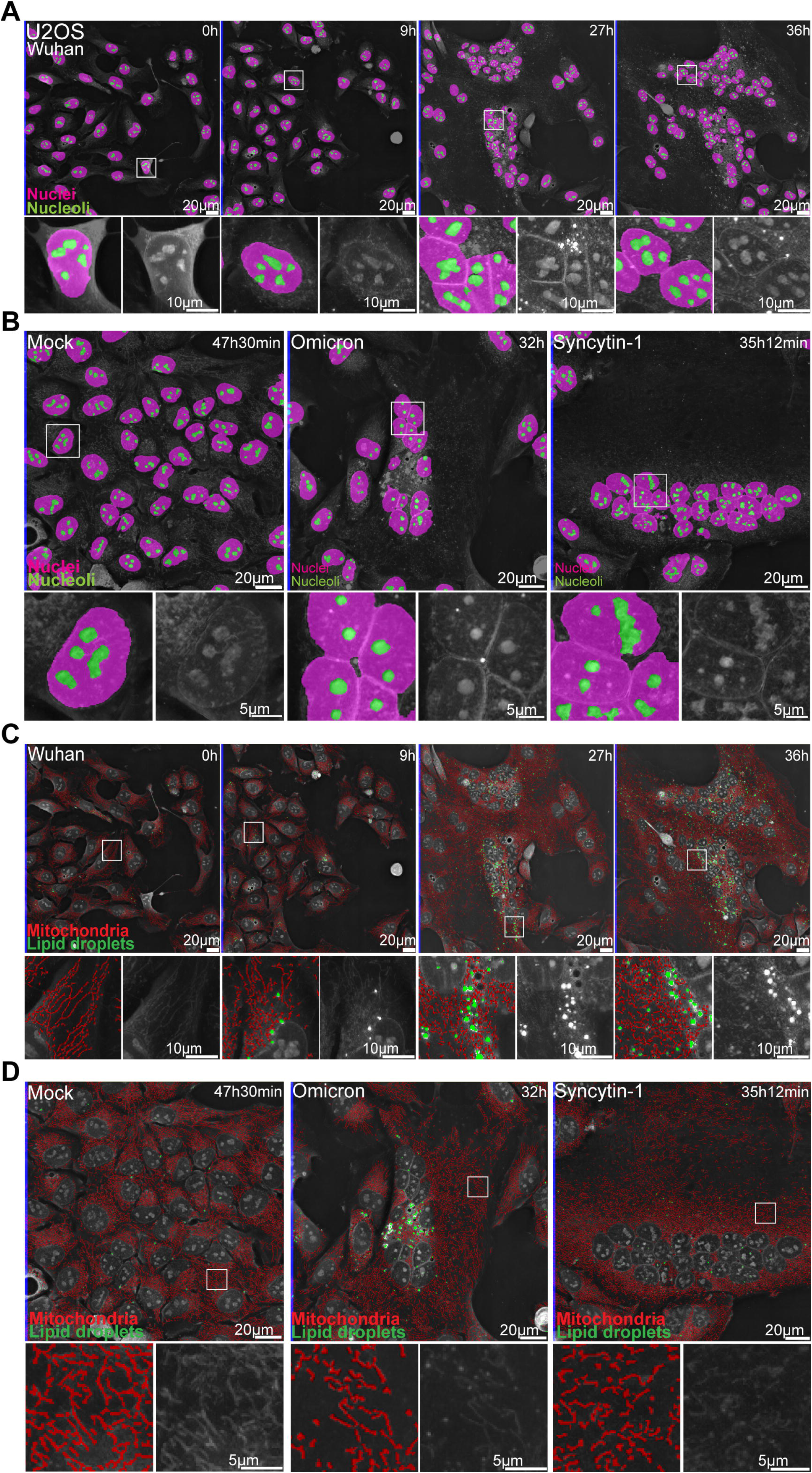
Automated detections of cellular organelles capture dynamics of SARS-CoV-2- induced changes. **(A)** 36 hours-long holotomographic microscopy (HTM) time lapse acquisition of U2OS cells infected by the Wuhan SARS-CoV-2 strain. Pink: nuclei. Green: nucleoli. **(B)** Late time point images of time lapse imaging experiments of non-infected cells (Mock), Omicron-infected cells and Syncytin-1-expressing cells. Pink: nuclei. Green: nucleoli. **(C)** 36 hours-long holotomographic microscopy (HTM) time lapse acquisition of U2OS cells infected by the Wuhan SARS-CoV-2 strain. Red: mitochondria. Green: lipid droplet. **(D)** Late time point images of time lapse imaging experiments of non-infected cells (Mock), Omicron-infected cells and Syncytin-1-expressing cells. Red: mitochondria. Green: lipid droplet.

We represented the data as time series of violin plots based on single cells or organelles points, to visualize the evolution of the various parameters over time and to allow explicit statistical assessment of infection-induced changes (Figure 4). The progression of nucleoli/nucleus size ratio was similar in Wuhan-infected and non-infected cells, as well as in syncytia triggered by Syncytin-1 (Figure 4A). In agreement with a recent report^39^, the nucleoli of Omicron-infected cells became larger, especially at a late time point of infection. This might reflect a recruitment of the nucleolar machinery to facilitate viral translation^40,41^. A massive increase in LD number and size was observed in SARS-CoV-2 Wuhan- or Omicron-infected cells but not in control or Syncytin-1-expressing cells (Figure 4B). These observations are in line with a SARS-CoV-2-induced remodeling of lipid metabolism^20^ to support pro-virus signaling^22^ and provide material for the formation of double membrane vesicles (DMV) (Figure 4B).

**Figure 4.**
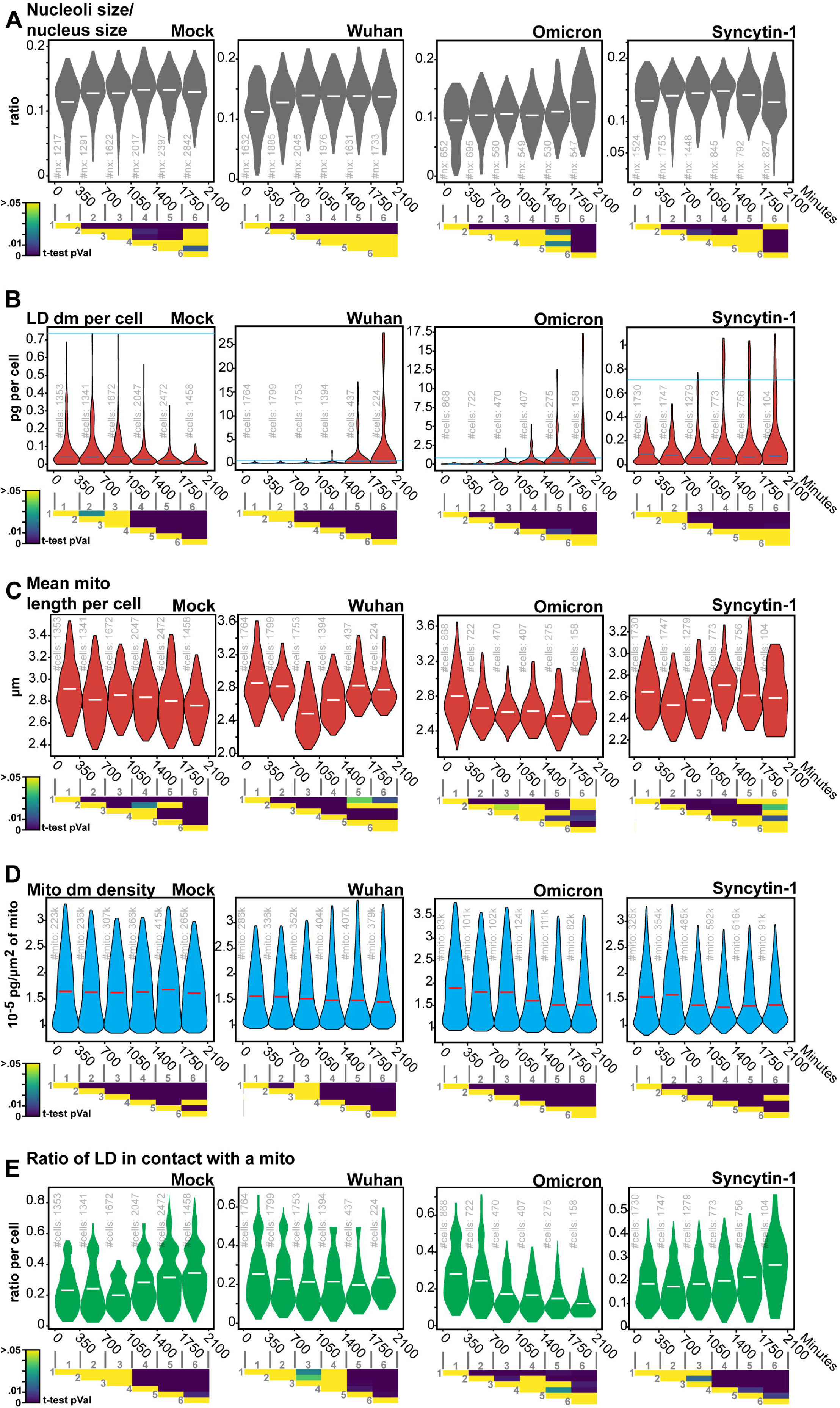
Label-free quantifications of SARS-CoV-2-induced alterations of organelle dynamics. **(A)**Nucleoli/nuclei size-ratio per cell over time (#of quantified nuclei in light-gray), **(B)** Lipid droplet dry mass per cell (#of quantified single cells in light-gray), **(C)** Mitochondria dry mass density (#of quantified mitochondria in light-gray), **(D)** Mean mitochondria length per cell (#of quantified cells in light-gray), **(E)** the ratio per cell of lipid droplets less than 400nm away from the nearest mitochondria (#of quantified cells in light-gray). Violin plot representation of non-infected, Wuhan-, Omicron-infected and Syncytin-1- expressing cells. Each violin plot represents the distribution of the segmented single cells or organelles contained within the indicated period. Bin-to-bin t-tests p-values are indicated below each experiment. Single cells and organelles studied in each of the mock, Wuhan, Omicron and Syncytin-1 are coming from at least 3 different movies (see Videos S1-S19).

The obvious effect of SARS-CoV-2 on LD indicated a broad metabolic impact and thus led to the question of mitochondrial alterations. In control cells, we detected a modest yet significant reduction of mitochondrial size over the 1.5 days of culture, which may reflect the changing metabolic state of proliferating cells that progressively consumed culture medium (Figure 4C). In both Wuhan and Omicron infected cells, there was a marked mitochondria size decrease around 15 h (900 min) pi (Figure 4C), that is likely an infection-induced imbalance of mitochondrial dynamics^42^. At later points, the length of the mitochondria increased again in infected cells. Contradictory findings of impaired mitochondrial fission and fusion in response to SARS-CoV-2 infection have been reported^43,44,45^.Our results suggest that these events may be temporally distinct.

We then quantified the dry mass, defined as the bulk content in biomolecules (mainly proteins, lipids and nucleic acid) that are not water of hundreds of thousands of mitochondria. In fact, HTM returns the refractive index of the observed biological structures, which is linearly linked to the content in biomolecules of the observed structure. In control cells, we observed a stable dry mass per unit of mitochondria size overtime (Figure 4D). This dynamic was changed in SARS-CoV-2-infected or Syncytin-1-expressing syncytia, which were characterized by an overall slight reduction of the dry mass of single mitochondria. This is in line with previous reports^45,46^ suggesting that SARS-CoV-2 down-regulates the translation of mitochondrial genes. The observation that Syncytin-1-expressing cells also displayed a decreased mitochondrial dry mass suggests that SARS-CoV-2 could use syncytia formation to promote mitochondrial alterations more efficiently.

We next took advantage of our capacity to localize organelles in space and relative to each other’s to measure the distance between LD and mitochondria, a marker of the rate of fatty acid oxidation and thus of energy production^33,34^. In uninfected or Syncytin-1- expressing cells, the proportion of LD in proximity (< 400 nm) of mitochondria increased over time (Figure 4E). In contrast, this ratio stayed stable or even decreased in Wuhan and Omicron-infected cells. Therefore, infection separates LDs from mitochondria, reflecting a probable impact of SARS-CoV-2 on cell metabolism.

### Large lipid droplets form in infected cells only

We next examined the links that may exist between LD, mitochondria, and viral production zones, that we identified by immuno-staining with anti-NSP3 or anti-double stranded (ds) RNA antibodies. The viral NSP3 protein plays many roles in the virus life cycle, including viral polyprotein processing, formation of the viral replication compartment, viral RNAs trafficking and innate immunity antagonism^47^. Anti-dsRNA antibody selectively recognizes viral RNAs and do not detect cellular RNAs. NSP-3 and dsRNA rarely colocalized within cells (Figure 5). Correlative HTM and confocal microscopy shows that at 24 h pi, cells that accumulated large perinuclear LD were also positive for dsRNA and NSP3 stainings (Figure 5A and S3), confirming that the accumulation of large LD is a signature of SARS-CoV-2 infection. dsRNA accumulated next to LD, while NSP3 was excluded from them (Figure 5B and 5C). We performed correlative HTM and electron microscopy (CHEM) of non-infected or infected cells and focused on areas displaying large LD. The mitochondria surrounded large LD in control cells but not in SARS-CoV-2 infected cells (Figure 5D, 5E and S4). Altogether, the HTM quantitative analysis, combined with qualitative correlative microscopy indicate that LD alteration together with mitochondria relocation is a hallmark of effective virus production.

**Figure 5.**
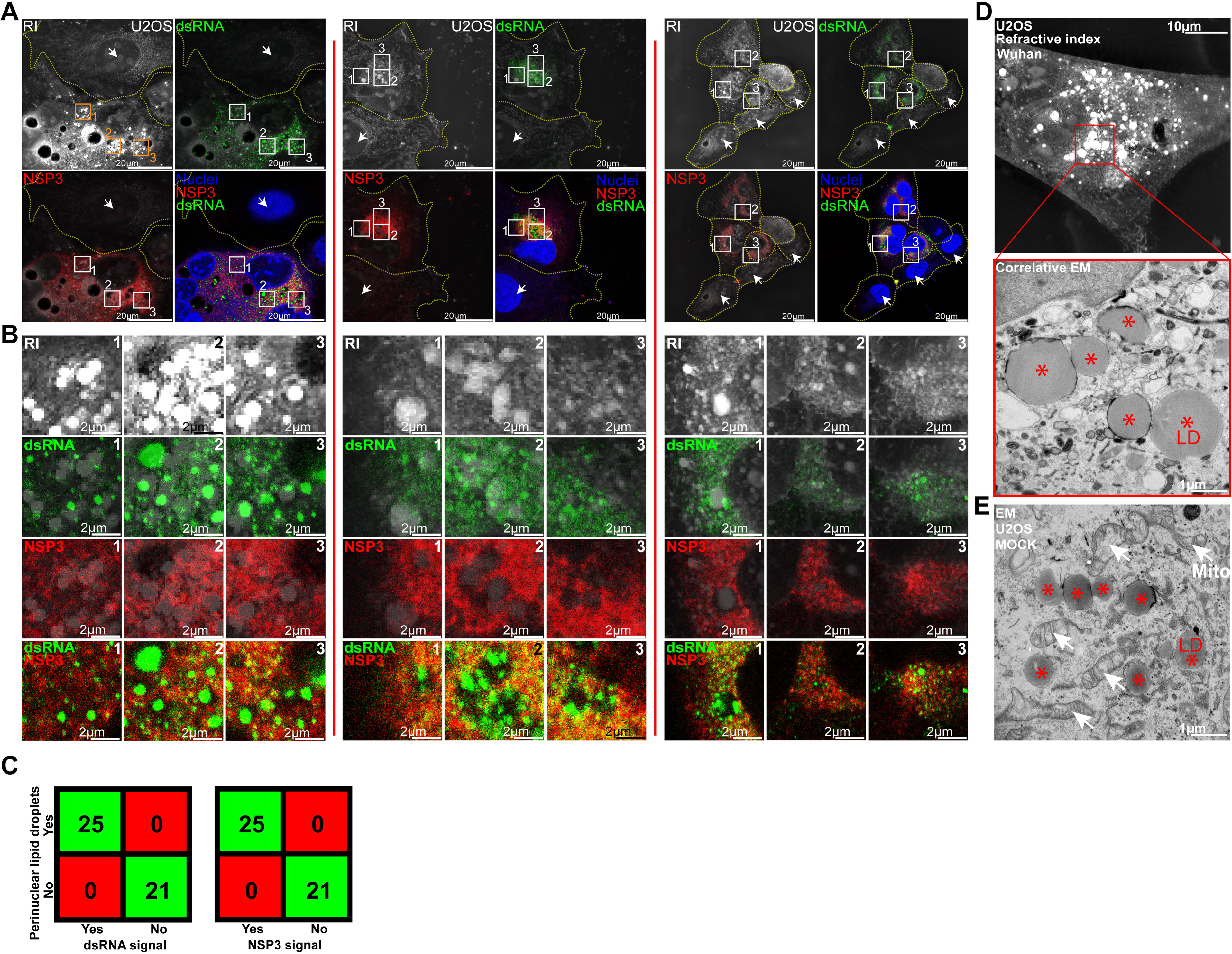
Perinuclear lipid droplet accumulation is a marker of infection. **(A-C)**Comparison of the refractive index (RI) signal acquired using holotomographic microscopy (HTM) with the double-stranded (ds) RNA and NSP3 immuno-fluorescent signals acquired with confocal microscopy. dsRNA spots are detected over and around lipid accumulations while the homogenous NSP3 signal is excluded from them. **(C)** Confusion matrix. Cells with perinuclear lipid droplet accumulation have a SARS-CoV-2-induced dsRNA and NSP3 signals. Cells with no lipid droplets (arrow) show no dsRNA and NSP3 signals. See supplementary S3. **(D)** Correlative holotomography/electron microscopy in infected cells. No mitochondria surround infection-induced lipid droplets. See also figure S4. **(E)** Electron microscopy in non-infected cells. Lipid droplets within are close to mitochondria (red stars indicate lipid droplets, white arrows indicate mitochondria).

### SARS-CoV-2 alters organelles cross-regulations

We next investigated the dependencies between LD, mitochondria, nuclei, nucleoli, and cytosol dry masses using comparative Bayesian networks (BN). BN are an established tool for modelling biological datasets^48^ in fields such as signaling^49^, genomics^50,51^, or immunology^52^, but not, to the best of our knowledge, to model the hierarchy and regulation existing between cell organelles. Established BN methods^53^ allow to search for the conditional relationships and probabilities between factors of interest. These methods provide an intuitive visual representation under the form of a directed acyclic graph. We thus investigated the likelihood of discretized organelles dry mass, given the dry mass of the other organelles. Considering that the dry mass variation of a subcellular compartment reflects regulated variations of its protein, lipid, and/or nucleic acid contents, we propose that the causal relationship between organelle dry mass captures an integrated level of organelle-dependent regulation. Henceforth, we will employ the term *organelle cross-regulation* (OCR) rather than referring to *dry mass influence diagrams*, the latter being less intuitive. As our approach relies on organelle detection, and not on gene or protein levels measurements, it is blind to organelle-independent global regulations that manifest during infection or cellular dysfunction, such as modulations of protein expression and other gene regulations that cannot be captured by assessing dry mass variations.

We also attributed a “regular” or “syncytium” identifying tag, to each cell of our control, SARS-COV-2-infected and Syncytin-1 expressing conditions. This allowed us to observe the specific impact of syncytia formation on OCR compared to infection. The differences between similarly established networks for the three conditions, provides unique insights on how SARS-CoV-2 infection or syncytia formation impacts OCR (Figure 6A-6C). The nucleus had an expected influence on nucleoli, irrespective of infection or syncytia formation. This indicates that our networks can capture relevant functional relationships between organelles. Moreover, in control cells, the network has in its center the nucleus dry mass. This was also expected, since in freely dividing cells, the cell cycle and thus the nucleus DNA content, must be central to all OCRs. We observed that SARS-CoV-2 infection rewired the OCR network, likely because it can alter organelles involved in numerous processes such as lipid metabolism^20–22^ and the cell cycle^54–57^. The main events driving the virus-induced OCR network were the formation of syncytia followed by the accumulation of LD (Figure 6B). This suggests that the SARS-CoV-2-induced syncytium has a role in promoting viral replication, apparently by boosting LD formation.

**Figure 6.**
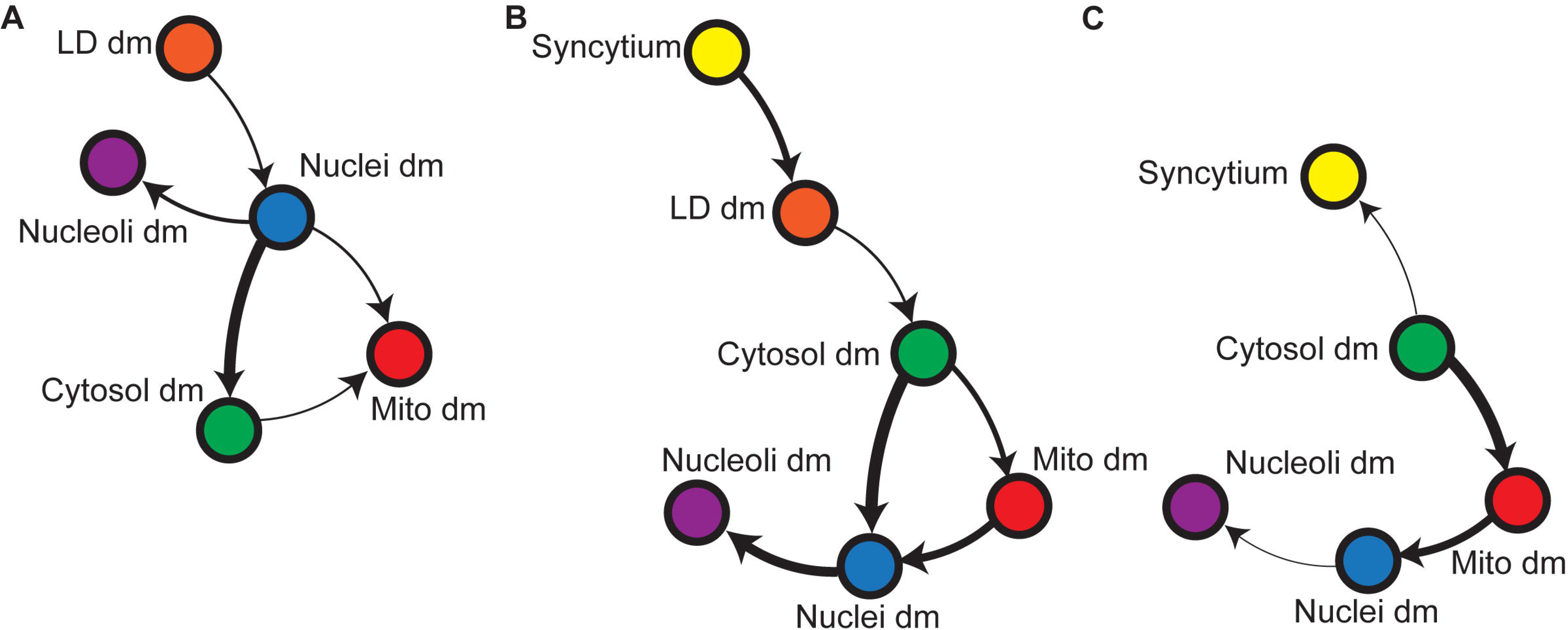
Comparative Bayesian networks show that SARS-CoV-2 infection changes organelle cross-regulation. **(A, B, C)**Bayesian networks representing the causal relationship between mitochondria, nuclei, nucleoli, lipid droplets and cytosol dry mass states and the syncytial state of **(A)** non-infected **cells (B)** infected cells **(C)** Syncytin-1expressing cells.

The formation of a syncytium is a complex, broad phenomenon that implies more than the plasma membrane fusion upon which we rely to identify it. The mechanisms at play to prepare its formation, cytoskeleton reorganization, nucleus clustering, or rapid mixing of different trafficking systems and signaling or metabolic states can be expected to have broad consequences that will not be captured as direct causal links but rather be seen in the way organelles are wired together. In cells forming syncytia independently of infection, the syncytial state had no direct impact on one specific organelle and was only loosely related to the rest of the network, with the cytosol dry mass as the sole, and faint, predictive factor (Figure 6C). We saw however the same hierarchy than in SARS-CoV-2 infected cells between the cytosol, the mitochondria, and the nuclei-nucleoli duo. Thus, the syncytium by itself changes how organelles interact together. This suggests that the virus may trigger the syncytium for its capacity to establish a broadly favorable OCR system. While the infectious and non-infectious syncytia looked similar in HTM images, the non-infectious syncytia lack the capacity to promote LD growth.

(Figure 4B and 6B). We used confocal microscopy of tubulin immuno-staining as a complementary approach to investigate further potential differences between the two types of syncytia. We observed that compared to Syncytin-1-induced syncytia, SARS-CoV-2-induced syncytia displayed a flat microtubule signal profile (Figure 7A and 7B) from the clustered nuclei towards the syncytium boundaries (Figure 7C). This flat microtubule signal distribution was detected in 77% of the infected syncytia and only in 20% of Syncytin-induced fused cells (Figure 7D). In the remaining 80% of Syncytin-induced fused cells, a gradient was seen, more tubulin was seen surrounding the nuclei cluster than in the cell boundaries. Altogether, these results indicate that in addition to the events captured by HTM, the cytoskeleton is differentially modified in SARS-CoV-2 infected or Syncytin1-induced syncytia, which could modulate LD accumulation and mitochondrial redistribution.

**Figure 7.**
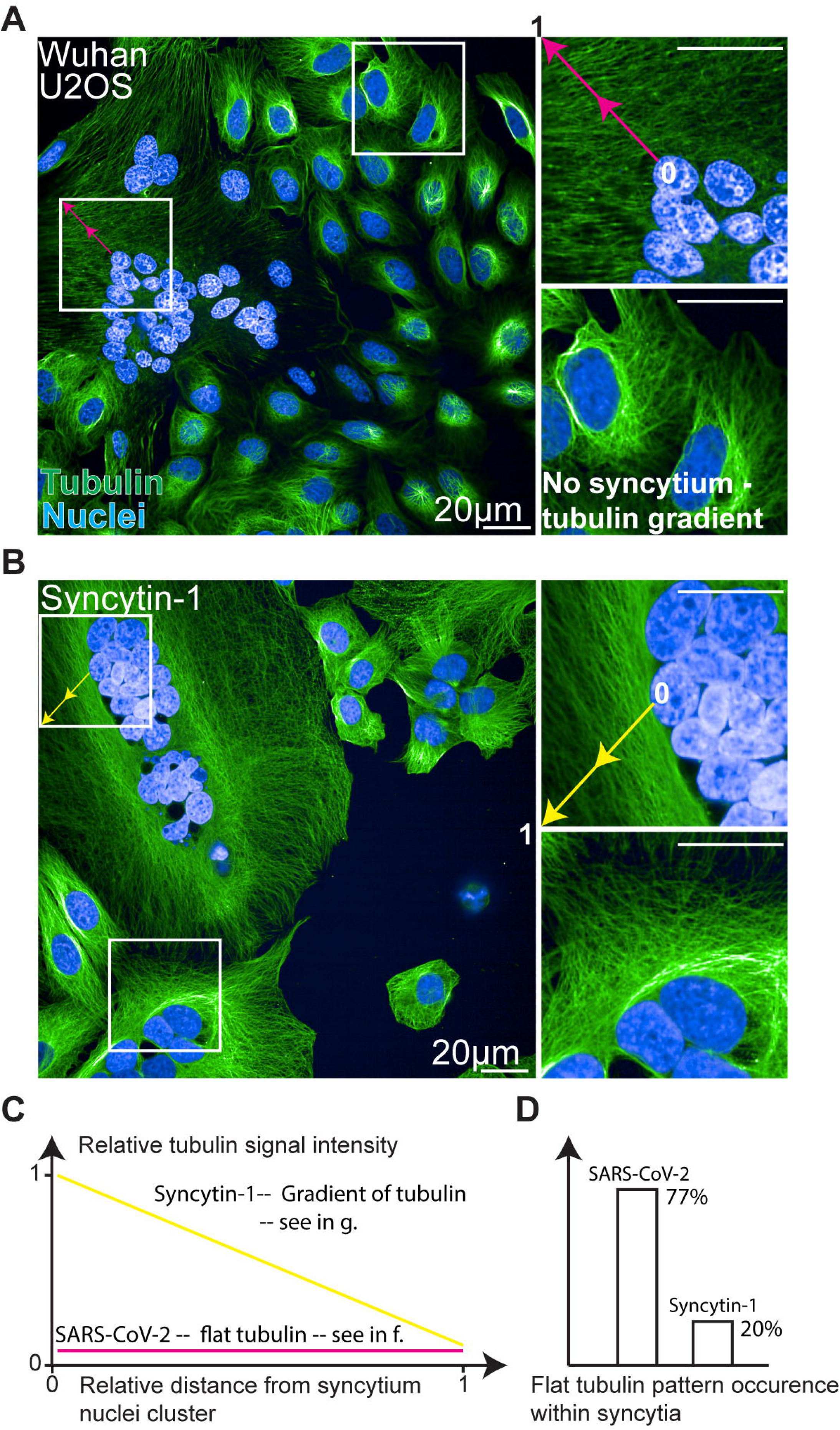
SARS-CoV-2-induced syncytia display flat radial tubulin signal. **(A,B)**Representative images of confocal immuno-fluorescence microscopy in SARS-CoV-2 Wuhan- or Syncytin-1- induced syncytia. Green: Tubulin. Blue: DAPI. **(C)** Representative tubulin signal profiles from a nuclei cluster towards cell boundaries **(D)** Occurrence of a flat radial actin pattern.

## Discussion

We demonstrate here that AI-enhanced label-free microscopy holds great promises for the quantitative investigation of pathological processes such as virus infection and offers new possibilities to quantitative cell biology in general^58^. The emergence of high-content fluorescence microscopy^59^ and massively multiplexed labeling methods^60^ revolutionized our understanding of cells’ systems^61^, whether they are unperturbed, adapting to changing micro-environments^62^ or reacting to drug treatments^63^. Yet, these impressive methods perturb living cells and are not suited for the monitoring of fine biological dynamics over long periods of time.

Amongst the currently available label-free microscopy techniques, holotomographic microscopy (HTM) produces images whose quality enable effective computer vision solutions^19^. To achieve this aim, it is important to overcome the laborious task of ground truth generation^64,65^ and to choose the adequate object detection technique, where deep learning is not always the most effective solution. The ground truth that we could produce for this work allowed for a U-NET segmentation of simple, large objects like cells or nucleoli, but fell short for a similar detection of mitochondria that are sparser, thinner objects. We speculate that the more explicit control of the receptive field in the case of the pixel classification approach, which depends on the blurring steps applied on images before feature extraction, was more adapted for objects spanning few pixels only. We believe that our work is particularly relevant to the field of quantitative mitochondrial biology^36,66,67^, given the susceptibility of that organelle to phototoxic stress and label-induced perturbations^8,68,69^.

Our object detection strategy transforms the limitation of HTM, which is the complex nature of the images it provides, into an advantage: having access to biologically relevant, direct, simultaneous measurements of organelles within single cells. This opens a vast landscape of possibilities for system’s investigations. Such dataset is particularity adapted for causal investigations due to the reasonable number of dependencies to evaluate. We could thus define in this study organelle cross-regulation (OCR) networks. There are pending questions: will the same OCR networks topologies be observed in other cell types or in response to other viruses? We believe that a comparative OCR analysis of different cellular conditions is essential. Future work will help determining whether we have uncovered conserved regulation networks for synchronizing organelles.

There are limitations to our study. Firstly, Bayesian networks are directed acyclic graphs and as such cannot be circular or contain feedback loops, which are natural components of many biological systems. Thus, the use of causal inference provides a partial view of OCR networks. Targeted metabolic or genetic perturbations to challenge the hypothesis gathered through network inference will help further characterizing OCR dependencies and their perturbations. Also, key OCR network dependencies could be further validated through HTM single-cell tracking. Secondly, key subcellular structures such as the Golgi or the endoplasmic reticulum are not captured by the current sensitivity of HTM and are therefore absent from our analysis. However, HTM can be associated with other label-free^70^, fluorescence or electron microscopy in correlative approaches. Such association is essential to advance both the label-free and label-based imaging worlds. This is illustrated here by the assessment of mitochondrial localization by CHEM, the colocalization of LD, dsRNA and NSP3 by correlative HTM/confocal microscopy, or the characterization of tubulin network modifications during SARS-CoV-2 infection and syncytia formation.

In this study, we used U2OS cells expressing the ACE2 receptor to establish a first set of discoveries in an uncharted territory. It will be important to extend such analysis to other cell types, for instance those that do not or poorly form syncytia upon infection. The use of viral strains carrying GFP-tagged viral proteins or other markers will facilitate the identification of infected cells^71^. This will allow to further explore the whole infection process and better characterize the role of syncytia during infection. Our work quantitatively describes the multiple cellular events associated with SARS-CoV-2 infection in real-time. An analysis of cells expressing individual viral proteins, such as for instance ORF9b^72^, that interacts with the mitochondrial protein TOMM70^73^, or NSP6, that mediates contact between DMVs and LDs^74^, will provide new clues on the impact of SARS-CoV-2 on the dynamics and shape of mitochondria.

In summary, we have developed a novel pipeline of analysis combing HTM, AI-assisted analysis and causality inferences using Bayesian statistics, to assess the impact of SARS-CoV-2 on the dynamics of cellular organelles and OCR. We report that the virus directly alters LDs, mitochondria, nuclei, and nucleoli as well as how those organelles influence each other. We also established that the infectious syncytium is likely favoring a pro-virus cellular environment. This approach opens exciting possibilities to analyze any pathogen, drug effects, and physiological or pathological events affecting the cell life, including nutrient variations, metabolic adaptation, and malignant transformation. It holds promise to lead to new insights into the dynamics a vast range of biological processes.

## Acknowledgments

We thank Timothée Bruel, Julian Buchrieser and Sasha Legrosdidier for critical reading of the manuscript, members of the Virus and Immunity Unit for discussions and help, Perrine Bomme and the Ultrastructural Bioimaging Unit (UBI) and UtechS Photonic BioImaging (UPBI) core facilities (Institut Pasteur). NS is funded by the ministère de l’Enseignement supérieur et de la Recherche. OS laboratory is funded by Institut Pasteur, Fondation pour la Recherche Médicale (FRM), ANRS-MIE, the Vaccine Research Institute (ANR-10-LABX- 77), HERA European program (DURABLE consortium), Labex IBEID (ANR-10-LABX-62- IBEID), ANR/FRM Flash Covid PROTEO-SARS-CoV-2 and IDISCOVR. We are grateful for support for equipment from the French Government Programme Investissements d’Avenir France BioImaging (FBI, N° ANR-10-INSB-04-01) and the French gouvernement (Agence Nationale de la Recherche) Investissement d’Avenir programme, Laboratoire d’Excellence “Integrative Biology of Emerging Infectious Diseases” (ANR-10-LABX-62-IBEID). We thank Thibault Corteoux and Lisa Polaro for connecting Nanolive’s Deep Quantitative Biology team with the Institut Pasteur. We thank all Nanolive’s employees for their encouragement and support.

## Author contributions

Conceptualization, N.S., B.M., T.W., O.S. and M.F.; Methodology, N.S., B.M., N.C., L.A., H.M., A.J., A.M., O.S. and M.F.; Data Curation, L.A., H.M., A.J., A.M. and M.F.; Software, L.A., H.M., A.J., A.M. and M.F.; Investigation, N.S., N.C., B.M. and M.F.; Visualisation, N.S. and M.F.; Writing – Original Draft, M.F; Writing – Review & Editing, N.S., T.W., O.S. and M.F.; Funding Acquisition, O.S.; Resources, O.S. and M.F.; Supervision, T.W., O.S. and M.F.

## Methods

### Holotomographic microscopy time lapse acquisitions

All label-free images were acquired using a 3D-Cell Explorer-fluo (Nanolive SA, Tolochenaz, Switzerland) microscope. This microscope is equipped with a 520nm laser for tomographic phase microscopy, the irradiance of the laser is 0,2nW/µm2 and the acquisition time per frame is 45ms. It is equipped with a Blaser ace acA1920-155um camera, and an air objective lens (NA = 0.8, magnification 60x). The microscope is equipped with a fluorescent module (pE-300ultra, CoolLED) for standard epifluorescence images of the sample in three different channels: Cy5 (excitation peak 635 nm), TritC (excitation peak 554 nm) and FitC (excitation peak 474nm). The microscope is equipped for long term live cell imaging: temperature, humidity, gas composition. The incubator chamber (Okolab) keeps the sample at 37°C, is closed by a heating glass lid to prevent condensation and is connected to a gas mixer (2GF-Mixer, Okolab) to maintain 5% of CO2. The humidity module ensures a 90% relative humidity within the chamber.

For imaging, 50’000 cells were plated in a 35mm No.1.5 ibidi polymer coverslip bottom dish. After 24 hours, the media was replaced with fresh media containing the indicated dose of virus. Dish were then placed in the incubator chamber of the microscope and imaging was started 3 hours post-infection.

### Live fluorescent controls

For mitochondria labeling, TMRE was added to the media for 30 minutes, before washing and cell imaging. For siRNA experiments, Dharmacon smartpool siRNA directed against human-OPA1 siRNA or a control siRNA was used. For nucleoli labeling, cells were transfected using lipo3000 as described by the manufacturer, with GFP-GRI, a subunit of Simian Foamy virus that contains a nucleolar localization signal fused with GFP^75^. 24 hours post-transfection cells were imaged using the Nanolive. For lipid droplet labeling, Bodipy 493/503 was added in serum free media for 15 minutes, before washing and imaging.

### Immunofluorescent labeling

Infected cells were fixed with 4% PFA for 30 minutes. For correlation with Nanolive microscopy, the dish was scrached to provide a visual landmark. They were then imaged on the Nanolive. They were permeabilized with 1% Triton for 10 minutes and blocked overnight with PBS/1% BSA/0.05% sodium azide. Mouse anti-dsRNA J2 (1:100, RNT-SCI-10010200; Jena Bioscience), sheep anti-SARS-CoV-2 nsp3 (1:200, DU67768, MRC PPU) were used overnight in blocking buffer. Donkey anti sheep 488 (#A-11015, ThermoFischer Scientific) and Donkey anti-mouse 647 (#A-31571, ThermoFischer Scientific) were used at a 1:500 dilution in blocking buffer for 1 hour. Imaging was performed using a Leica SP8 confocal microscope. For cytoskeleton imaging, infection was performed in a 96 well plate (Perkin Elmer). Infected cells were fixed with 4% PFA for 30 minutes, permeabilized with 1% Triton for 10 minutes and blocked overnight with PBS/1% BSA/0.05% sodium azide. Mouse anti-tubulin (1:100, ProteinTech 66031-1-Ig) was used overnight in blocking buffer. Goat anti-mouse 488 (#A-11015, ThermoFischer Scientific, # A-11001) was used at a 1:500 dilution in blocking buffer for 1 hour. Imaging was performed using the Operetta Phenix (Perkin Elmer).

### Electron microscopy

50’000 cells were plated in a MatTek 35 mm Dish (P35G-1.5-14-C-GRD). The next day, the media was replaced with fresh media containing the indicated dose of virus. After 24 hours, they were fixed with 4% PFA for 30 minutes, and imaged using holotomographic microscopy. Cells were then fixed in 2.5% Glutaraldehyde (Sigma) in 1X PHEMS buffer overnight at 4°C. Samples were washed 3 times in 1X PHEM buffer and post-fixed in 2% OsO_4_ (Electron Microscopy Sciences) +1,5% potassium ferrocyanide in water 1h in the dark. After 3 washes in water they were incubated 20min in 1% uranyl acetate in ethanol 25%. Samples were gradually dehydrated in an ethanol series (50%,75%,95%, 3×100%) and then embedded in EMbed-812 epoxy resin (Electron Microscopy Sciences), followed by polymerization for 48h at 60°C.Thin sections (70nm) of the region of interest were cut with a Leica Ultramicrotome Ulracut UCT stained with uranyl acetate and lead citrate. Images were acquired using a Tecnai T12 120 kV (Thermofisher) with bottom-monted EAGLE camera.

### Cells

HeLa, 3T3-derived preadipocytes, HEPG2 and U2OS cells were purchased from ATCC. U2OS cells were stably transduced with pLenti6-human-ACE2 as previously described^28^ or with pCW57.1_Syn1. PCW57.1_Syn1 was obtained by cloning the previously described phCMV_Syn1^76^ into pCW57.1 (Addgene) using the NheI+AgeI restriction for both plasmids. All cells were cultured in DMEM with 10% fetal bovine serum (FBS), 1% penicillin/streptomycin (PS). For imaging and infections 25nM HEPES was added to the media. 10 μg/ml blasticidin were added to U2OS cells cultures. Cells were routinely screened for mycoplasma.

### Virus

The strain BetaCoV/France/IDF0372/2020 was supplied by the National Reference Centre for Respiratory Viruses hosted by Institut Pasteur (Paris, France) and headed by Pr. S. van der Werf. The human sample from which the strain was isolated has been provided by Dr. X. Lescure and Pr. Y. Yazdanpanah from the Bichat Hospital, Paris, France. The viral strain was supplied through the European Virus Archive goes Global (Evag) platform, a project that has received funding from the European Union’s Horizon 2020 research and innovation program under grant agreement no. 653316. Titration of viral stocks was performed on Vero E6, with a limiting dilution technique allowing a calculation of DCP50.

### Holotomographic image processing

All the time points composing the time-lapse imaging experiments were reconstructed and stored as 3D volumes by Eve, the software that controls the 3D cell explorer microscope and early data management. Custom routines were used to convert 3D volumes into 2D maximal projections along the z-axis.

### Deep Learning object detection

A custom U-Net architecture was used to segment nuclei, nucleoli, and cells. Training processes were run for 500 epochs for the nuclei and nucleoli models on an Nvidia RTX3060 GPU with 12GB of VRAM and for 50 epochs for the cell models on an Nvidia RTX A4000 16GB of VRAM. The ground truths for the cells, nuclei, and nucleoli, were made semi-manually with the help of a custom labeling tool and the guidance of fluorescently stained cells.

### Nucleus

Our nucleoli model was trained with a dataset of 655 holotomographic microscopy images of mammalian cells images randomly split into a training set (589 images) and a testing set (66 images). These images come from 58 acquisitions and include multiple cell lines. Each image contains from 1 to 23 nuclei, all have a size of 480×480 pixel achieved by cropping or zero-padding. Probability maps provided by the model are binarize with a 50% probability threshold. All objects smaller than 10 pixels are rejected. The training took 4 hours and 19 minutes.

### Nucleoli

Our nucleoli model was trained with a dataset of 495 holotomographic microscopy images of mammalian cells randomly split into a training set (445 images) and a testing set (50 images). Each image contains at least a dozen of nucleoli and have a size of 480×480 pixel achieved by cropping or zero-padding. Probability maps provided by the model are binarized with a 50% probability threshold. All objects smaller than 10 pixels are rejected. Potential unspecific signal outside the nucleus is removed thanks to the nucleus masks predicted by our nucleus model. The training took 4 hours and 13 minutes.

### Cells segmentation

The first part of our cell segmentation process aims at obtaining a rough cell segmentation without a precise cell border detection. This U-Net model was trained with a dataset of 1445 holotomographic microscopy images of mammalian cells randomly split into a training set (1295 images) and a testing set (150 images). Each image contains from 1 to 10 cells and have a size of 480×480 pixel achieved by cropping or zero-padding. Probability maps provided by the model are binarized with a 50% probability threshold. All objects smaller than 10 pixels are rejected. The training took 51 minutes. The second part of our cell segmentation process takes the U-Net produced cell blobs binary masks as a seed for a precise cell edges detection using a propagation algorithm approach.

### Ensemble pixel classification for mitochondria detection

Mitochondria were segmented using a tailored pixel classification approach inspired by previous seminal work^31^. Because the simple refractive index value is not a reliable organelle signature, we increased the pixel feature space dimensionality by applying on refractive index images a set of convolution filters using the VIGRA Computer Vision Library (v1.11.1) for a total of 21 dimensions. Thus, each pixel is described by its respective refractive index value and by its value in the 20 convolved images created by applying the following 10 filters on the refractive index image at two different sigma values (1.4 and 2.0): gaussian smoothing, difference of gaussian, gaussian gradient, gaussian gradient magnitude, hessian of gaussian, hessian of gaussian eigenvalue, Laplacian of gaussian, structure tensor eigenvalue, tensor determinant, and tensor trace. The probability for each augmented pixel of our images to be part of a mitochondria or lipid droplet was then evaluated by an extra-tree classifier (scikit-learn v1.2.2) diverging from the default hyperparameter setup only by the number of estimators that is equal to 200. Each resulting probability image is then transformed into a binary mask using an adaptive background removal approach (OpenCV, cv2.threshold). Our mitochondria extra-tree classifier was trained thanks to the labeling of 149 images from 2 different cell lines (CHO and Preadipocytes) coming from 14 different acquisitions.

### Detection of lipid droplets

Lipid droplets were automatically segmented using Nanolive lipid droplet assay software that identifies spherical structures of even signal distribution over specific local refractive index maxima. The produced lipid droplet segmentations were further used for custom metrics calculations.

### Metrics calculation

The instantiated masks of cells, mitochondria, lipid droplets, nuclei and nucleoli and cytosol were used to extract the spatial (shape and size descriptors, centroid position) and RI-derived (dry mass, textures) features of each of the mentioned biological objects in each frame of each time-lapse experiment (Video S1-S19). This was performed using a scientific python environment and the library scikit-image. Each object’s dry mass content was calculated from its refractive index value using linear calibration model [32]. The data were exported as a .csv file into a Python environment and plotted with the Python library matplotlib for figure making or used through the bnlearn python library to establish our organelle cross-regulation networks.

### Bayesian network structure learning

We performed our Bayesian network (BN) structure learning using the bnlearn^53^ python package, to determine which BN captures the best the causal links that exists between our dataset variables, lipid droplets, mitochondria, nucleus, nucleoli and cytosol dry mass densities, as well as the nature of the single cell, normal or syncytium.

The best BN for each condition was defined using a greedy score-based structure learning strategy. For this study we used the K2 scoring function and the hill climb search algorithm, which incrementally tests BN alternatives in order to improve the K2 scoring that itself determines the probability of the BN structure given its training data.

The independence test to define edge strengths of our BN was a chi squared statistical test using each of the three models and their related data. For each pair of our learned BN, a chi squared statistical test is performed to determine its pValue and associated chi squared score. The latter defines the thickness of the displayed edges of our organelle cross regulation Bayesian networks in Figure 6.

## Supplemental information titles and legends

**Figure S1.**
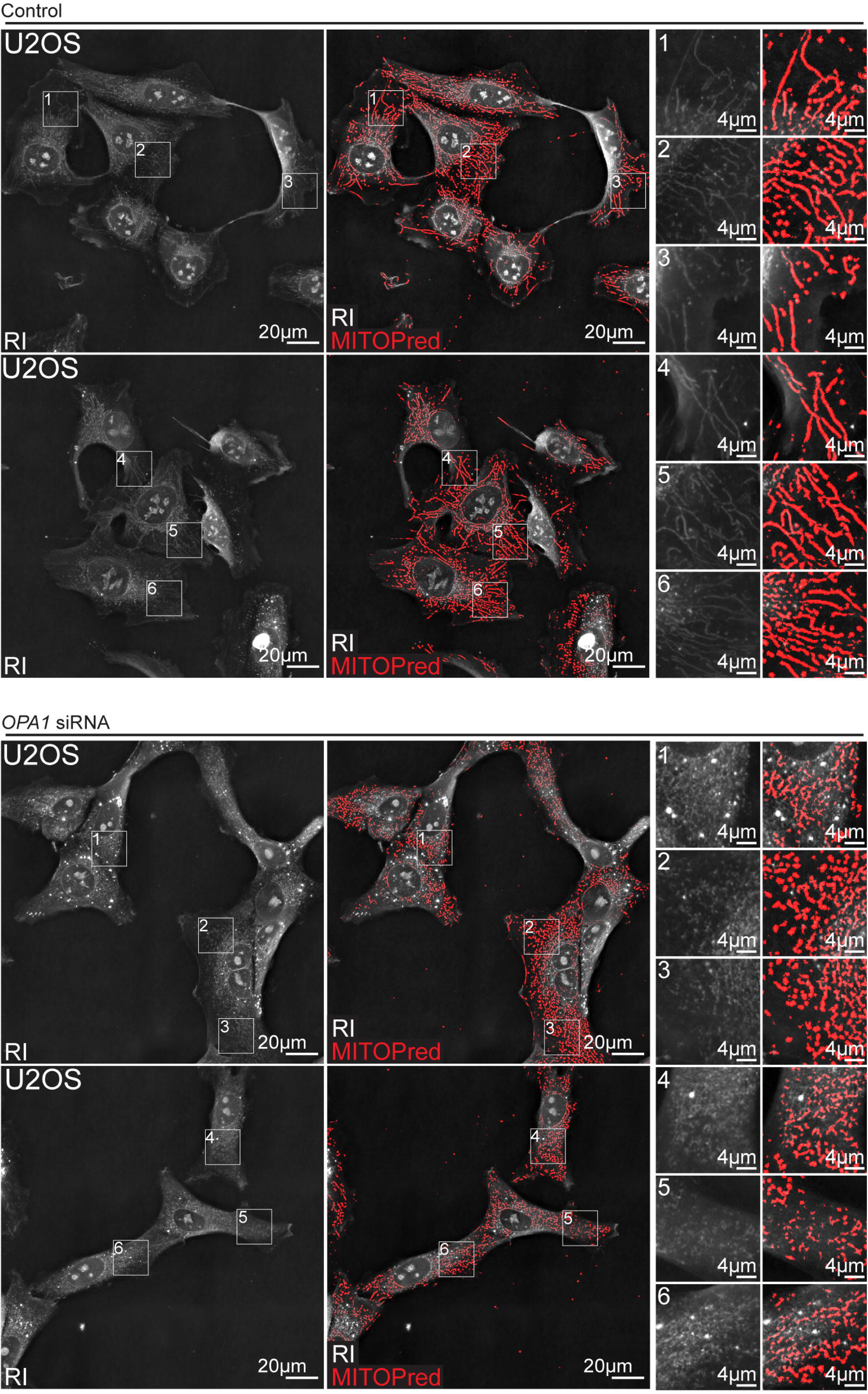
Machine learning detects unperturbed or fragmented mitochondria after silencing of *OPA1*.

**Figure S2.**
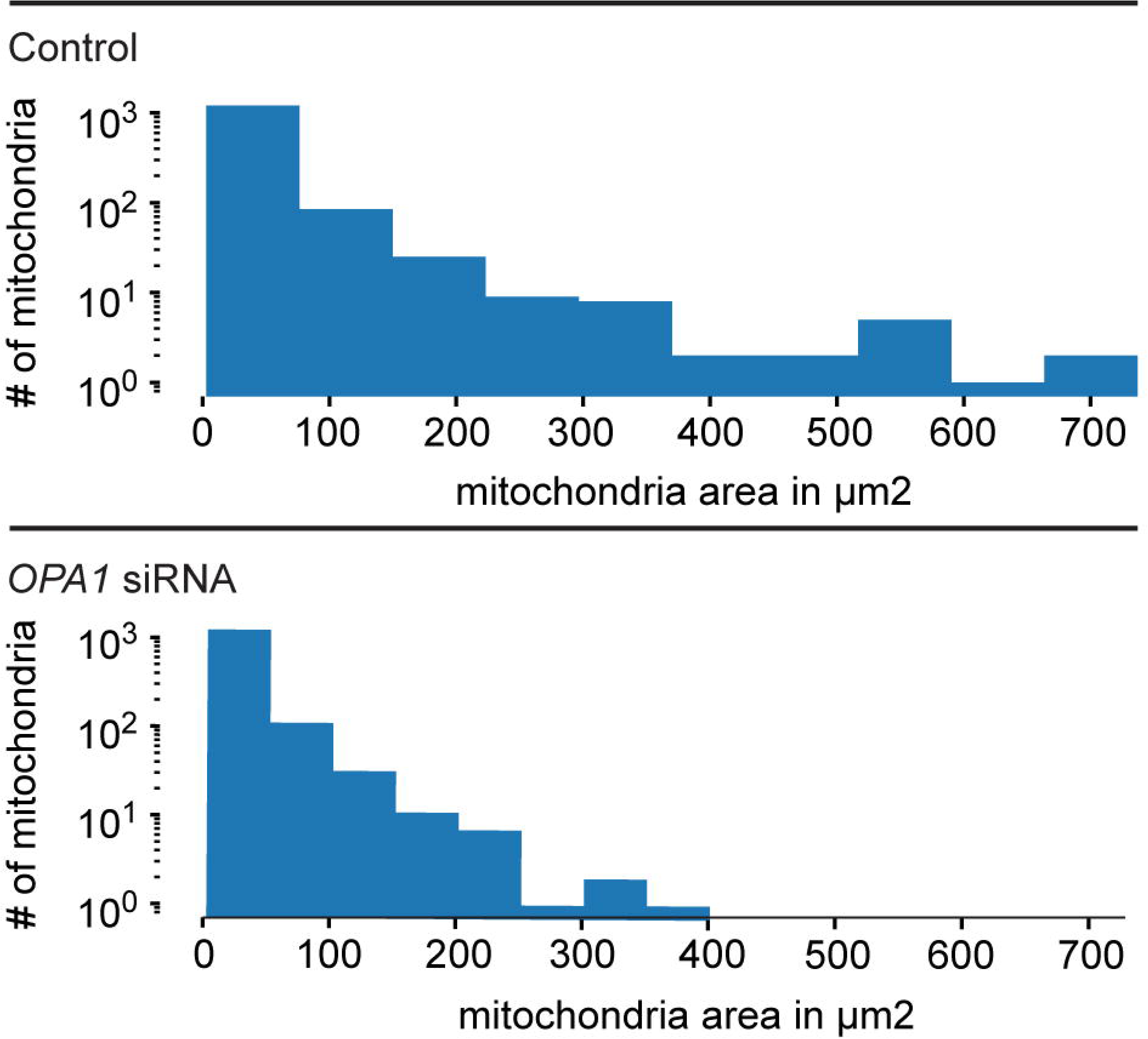
The effect of *OPA1* silencing is quantified through automated mitochondria segmentation.

**Figure S3.**
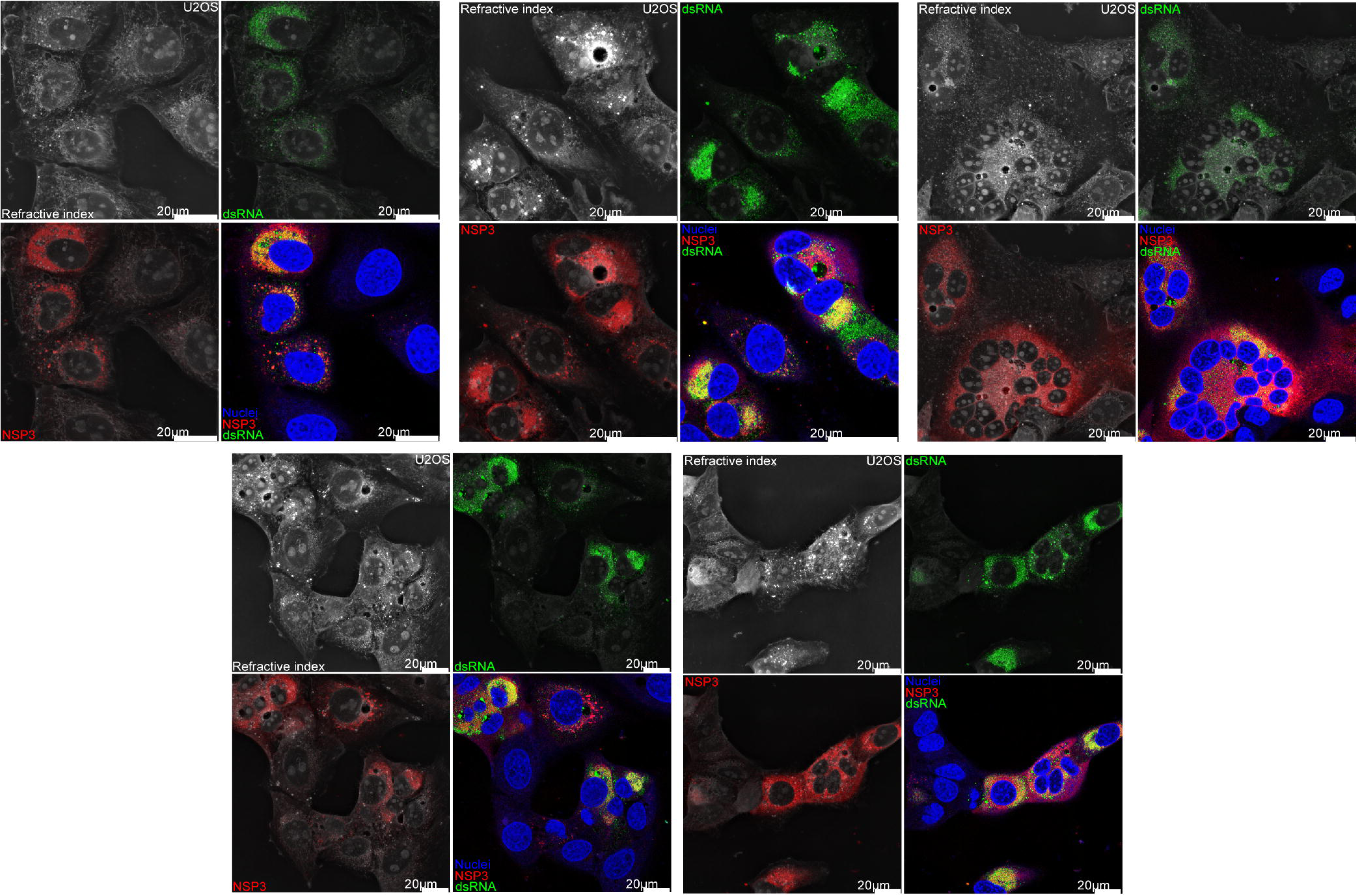
Perinuclear lipid droplet accumulation is a marker of infection.

**Figure S4.**
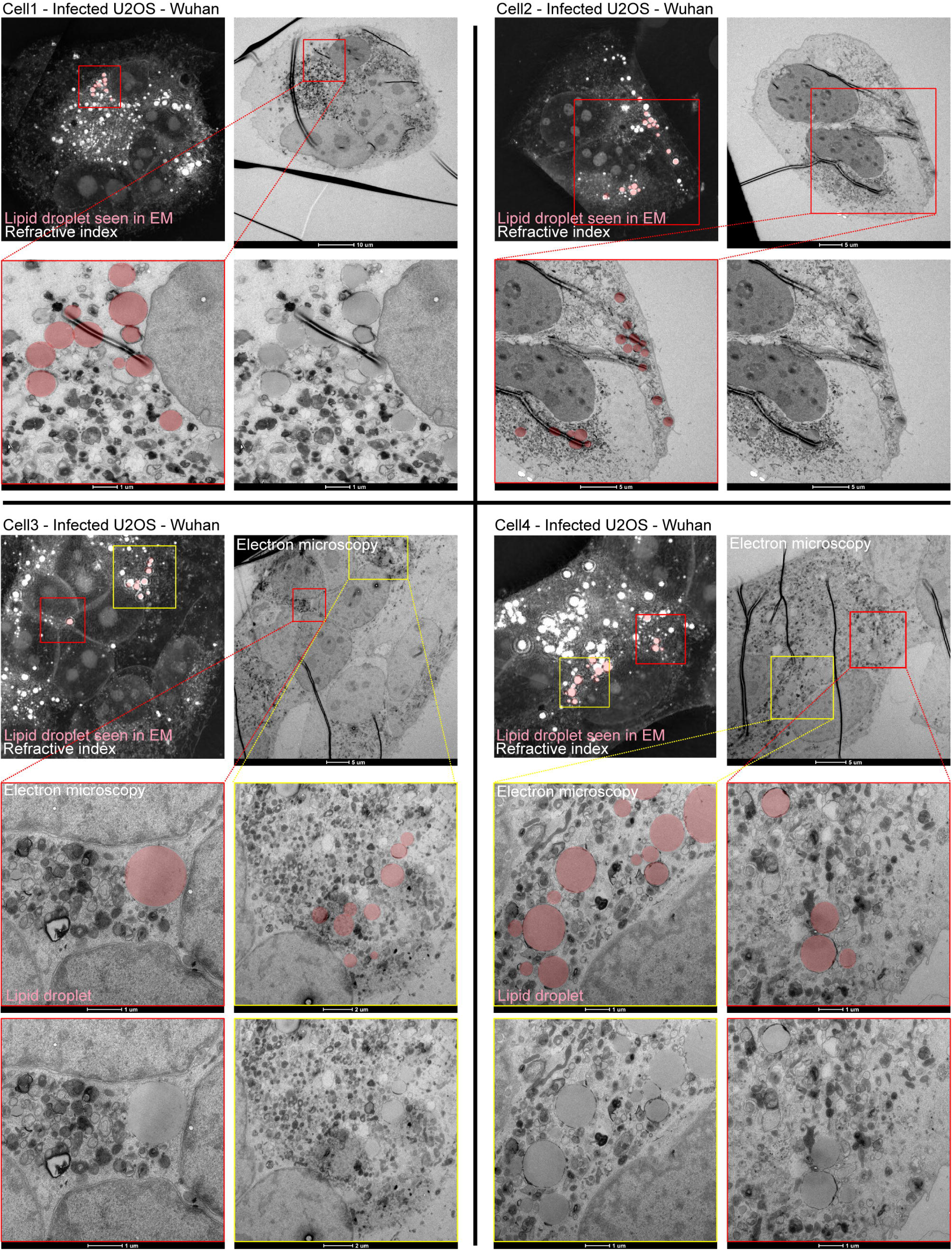
Lipid droplets of infected cells are not surrounded by mitochondria.

**Videos S1-S7: Control uninfected U2OS-ACE2 cells. Seven representative videos of cells and organelles masks overlaid on refractive index signal.**

**Videos S8-S12. U2OS-ACE2 cells infected with SARS-CoV-2 Wuhan strain. - Five representative videos of cells and organelles masks overlaid on refractive index signal.**

**Videos S13-S16. U2OS-ACE2 cells infected with SARS-CoV-2 Omicron BA.1. - Four representative movies of cells and organelles masks overlaid on refractive index signal.**

**Videos S17-S19. U2OS-ACE2 cells forming syncytia upon expression of Syncytin-1. - Three representative videos of cells and organelles masks overlaid on refractive index signal.**

